# Flow cytometry-based determination of ploidy from dried leaf specimens in genomically complex collections of the tropical forage grass *Urochloa* s. l

**DOI:** 10.1101/2021.03.26.437252

**Authors:** Paulina Tomaszewska, Till K. Pellny, Luis Miguel Hernández, Rowan A. C. Mitchell, Valheria Castiblanco, José J. de Vega, Trude Schwarzacher, Pat (J.S.) Heslop-Harrison

## Abstract

We aimed to develop an optimized approach to determine ploidy for dried leaf material in a germplasm collection of a tropical forage grass group, including approaches to collect, dry and preserve plant samples for flow cytometry analysis. *Urochloa* (including *Brachiaria, Megathyrus* and some *Panicum*) tropical grasses are native to Africa and are now, after selection and breeding, planted worldwide, particularly in South America, as important forages with huge potential for further sustainable improvement and conservation of grasslands. The methods enable robust identification of ploidy levels (coefficient of variation, CV, typically <5%). Ploidy of some 353 forage grass accessions (ploidy range from 2 to 9), from international genetic resource collections, showing variation in basic chromosome numbers and reproduction modes (apomixis and sexual), were determined using our defined standard protocol. Two major *Urochloa* agamic complexes used in the current breeding programs at CIAT and EMBRAPA: the ‘*brizantha*’ and ‘*humidicola*’ agamic complexes are variable, with multiple ploidy levels and DNA content. *U. brizantha* has odd level of ploidy (*x*=5), and the relative differences in nuclear DNA content between adjacent cytotypes is reduced, thus more precise examination of this species is required. Ploidy measurement of *U. humidicola* revealed some aneuploidy.

## 1. Introduction

Understanding the genome compositions of species within complexes including diploid and polyploid species is critical to evaluate their biodiversity, for conservation and to evaluate the potential for use in breeding; measurement of genome size, across potentially large germplasm collections, underpins such work. Grasslands and rangelands with grasses as the dominant species, being the largest ecosystems in the world, are the basic feed resources for livestock, and contribute to the livelihoods of over 800 million people including smallholders (Food and Agriculture Organization of the United Nations; http://www.fao.org/). Only 100–150 of the 10,000 forage species have been extensively cultivated, but many more have great potential for sustainable agriculture, and improvement and conservation of grasslands, including the genus *Urochloa* (previously classified in *Brachiaria*, and some *Eriochloa, Panicum* and *Megathyrsus*; González and Morton, 2005) comprising species native to tropical and subtropical regions of Africa. The great forage potential of these grasses have been recognized in the 1950s (Miles *et al*., 1996), leading to the acquisition of 700 accessions of *Urochloa* and related genera during the joint collection mission of CGIAR (Consultative Group on International Agricultural Research) lead centers: CIAT (Centro Internacional de Agricultura Tropical) and ILRI (International Livestock Research Institute) in Africa in the 1980s. Five species of *Urochloa*: *U. ruziziensis, U. decumbens, U. brizantha, U. humidicola*, and *U. maxima* were then introduced in South America, and have been using as fodder plants mainly in Colombia and Brazil (Keller-Grein *et al*., 1996).

In exploiting biodiversity in breeding, improvements in yield and nutritional quality of forages can be achieved by identifying genes increasing the digestibility of plant cell walls and the protein and lipid content in vegetative tissues, and increasing biomass production (Capstaff and Miller, 2018). By introduction to plant breeding programmes, genetic improvement of forage lines, recurrent genetic selection of plants showing useful traits, and subsequent hybridizations and back-crossings (Barrios *et al*., 2013; Hanley *et al*., 2020), create more diverse agroecosystems resilient to climate and environmental changes (Baptistella *et al*., 2020). The DNA amount measurement for ploidy and genome size estimation, and the characterization of genome composition are required for effective use of diploids and polyploids in breeding programs, as well as for research purposes (Ochatt, 2008; Tomaszewska *et al*., 2021).

Preparation of metaphases from dividing plant tissues, followed by microscopy and chromosome counting, is widely used to determine the ploidy of individual plants and show polyploid series within larger groups. However, the method is time-consuming and highly skilled, both in terms of growing plants and collecting root-tips or meiotic material, and in making the preparations. The most rapid and convenient technique for ploidy measurement is flow cytometry using suspensions of fluorescently labelled nuclei (Heslop-Harrison and Schwarzacher, 1996; Doležel *et al*., 1997; Schwarzacher *et al*., 1997; Bennett *et al*., 2000; Śliwińska, 2018), that is now widely adopted for fresh leaf specimens. Image cytometry is another tool for nuclear genome size analysis; however, despite some examples of its use (Greilhuber *et al*., 2003), it has not proven widely applicable for estimation of ploidy in plant tissues, because its image processing algorithms gives imprecise and unreliable results that cannot be compared to other methods (Svoboda *et al*., 2009). Scanning microdensitometry has proved reliable in measuring genome sizes, but requires equipment not now available (Bory *et al*., 2008). Direct sequencing of DNA, and either assembly or analysis of counts of short sequence motifs present in the reads (k-mer analysis) is often used to measure genome sizes in DNA sequencing programmes (Marçais and Kingsford, 2011), but is not appropriate for screening collections.

Phenolics, hydroxamic acids, and short-chain fatty acids are present in plants, and some of these phytochemicals have been identified as inhibitors of fluorescent DNA staining, hence leading to inaccurate flow cytometry-based measurement of DNA content (Loureiro *et al*., 2006a; Bennett *et al*., 2008; Greilhuber, 2008; Price *et al*., 2000; Jędrzejczyk and Śliwińska, 2010). The ability of tropical and subtropical plants to synthesize secondary metabolites and possess allelopathic potential is exceptional (Ooka and Owens, 2018). Mild winters and small temperature fluctuations mean that the growing season is year-round in tropical and subtropical regions, and they provide the strong competition of plants for resources, and succession. Seasonal and regional differences in accumulation of secondary products may cause differences in staining for flow cytometry. Secondary metabolites and their phytotoxicity on forage legumes have been recognized in *Urochloa* tropical forage grasses (Ribeiro *et al*., 2012, 2018; Oliveira *et al*., 2017; Feitoza *et al*., 2020), which has been suggested to make it difficult to analyze these plants by flow cytometry (Penteado *et al*., 2000).

For *Urochloa*, ploidy estimation across the whole germplasm collection (excluding one species, *U. ruziziensis*, known only as a diploid) is required due to the different pathways of reproduction showing sexual and apomictic accessions within same species (Roche *et al*., 2001), natural triploid interspecific hybrids (Timbó *et al*., 2014), different genome compositions both within and between species (Tomaszewska *et al*., 2021), confirmed aneuploidy (Moraes *et al*., 2019; da Rocha *et al*., 2019; Tomaszewska *et al*., 2021), and different basic chromosome numbers (*x*=6, 7, 8 and 9; de Wet, 1986; Basappa *et al*., 1987; Bernini and Marin-Morales, 2001; Risso-Pascotto *et al*., 2006; Boldrini *et al*., 2009b; Worthington *et al*., 2019).

Ideally, a common reference standard for flow cytometry and ploidy measurement should be a diploid plant from the taxon of the tested samples, grown and collected under similar conditions, and where chromosomes can be prepared and counted. For a pool with diverse ploidies, several standards are helpful, although the lack of seeds or living plants may make it impossible to prepare metaphase plates from root tips, and challenges (e.g. due to apomictic mode of reproduction of studied species, difficulties with germination of tropical plant seeds, or having only herbarium samples), may mean a less related standard with known ploidy level, genome size and basic chromosome number similar to the unknown samples must be used (Śliwińska, 2018).

Fresh leaves usually been considered the best material for flow cytometry analysis. However, there is often a requirement for use of field-material, collected under sub-optimal demanding conditions compared to plants for greenhouse or experimental field and requiring storage and transport to the flow cytometry facility. Also, work often needs to use herbarium or stored material, which may not be possible to collect again, or is the reference for published studies, or is determined as a new species/taxonomic revision, requiring determination of ploidy and estimation of genome size (Suda and Trávníček, 2006). The applicability of flow cytometry for dehydrated leaves is limited by several factors, including insufficient amounts of tissue, sampling of mature plants, incorrect drying, storage and preservation of samples, and the low efficiency of nuclei isolation due to their degradation.

For flow cytometry analysis of nuclear genome sizes from fresh and dried material, coefficient of variation (CV) – defining the variation in fluorescence intensity from nucleus to nucleus, visualized as the width of the intensity peak – is an important criterion showing estimation of nuclei integrity and variation in DNA staining (Bennett and Smith, 1976; Bennett *et al*., 1982; Arumuganathan and Earle, 1991; Bennett and Leitch, 2011; Loureiro *et al*., 2006b). Low coefficient of variation, even for dried leaf specimens (Suda and Trávníček, 2006), or seeds (Jędrzejczyk and Śliwińska, 2010), petals and pollen (Roberts, 2007), can be achieved by using appropriate isolation buffers and their supplementation, stains and staining protocols, the practical technique used for chopping leaves in the buffer, and even choice of razor blades.

Here, we aimed to develop an optimized and robust approach to determine ploidy for dried leaf material of tropical forage grasses. The established method can be widely adopted for dried leaf specimens, especially when screening genomically variable germplasm resource collections, defining a standard protocol recommendation. More specifically, we intended to optimize the flow cytometry assay for *Urochloa* grass group that shows variation in basic chromosome numbers and reproduction modes.

## 2. Materials and Methods

### 2.1. Plant material

Accessions of *Urochloa* and related species used in the study are listed in **Supplementary Data Table S1**. Seeds of *Urochloa* sp. PI 657653, *U. brizantha* PI 292187 and *U. maxima* PI 284156 (**Table 1**) were provided by United States Department of Agriculture (USDA, USA). Seeds of *Panicum miliaceum* Mil69 were provided by the Vavilov Research Institute (VIR, St Petersburg, Russia). Centro Internacional de Agricultura Tropical (CIAT, Colombia) provided seed samples of *U. decumbens* 664 and 6370, *U. ruziziensis* 6419, *U. humidicola* 26151 and 16867, and *U. maxima* 6171 and 16004. Leaf samples intended for flow cytometry analysis were collected from germplasm accessions grown and maintained in the field genebank at CIAT Palmira campus (**Fig. 1**).

**Table 1.**
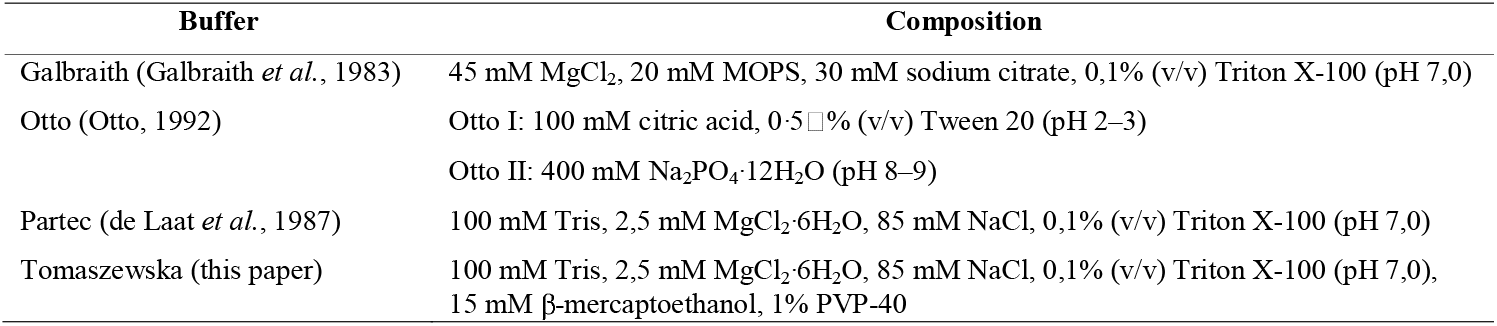
Nuclei isolation buffers and their compositions.

**Figure 1.**
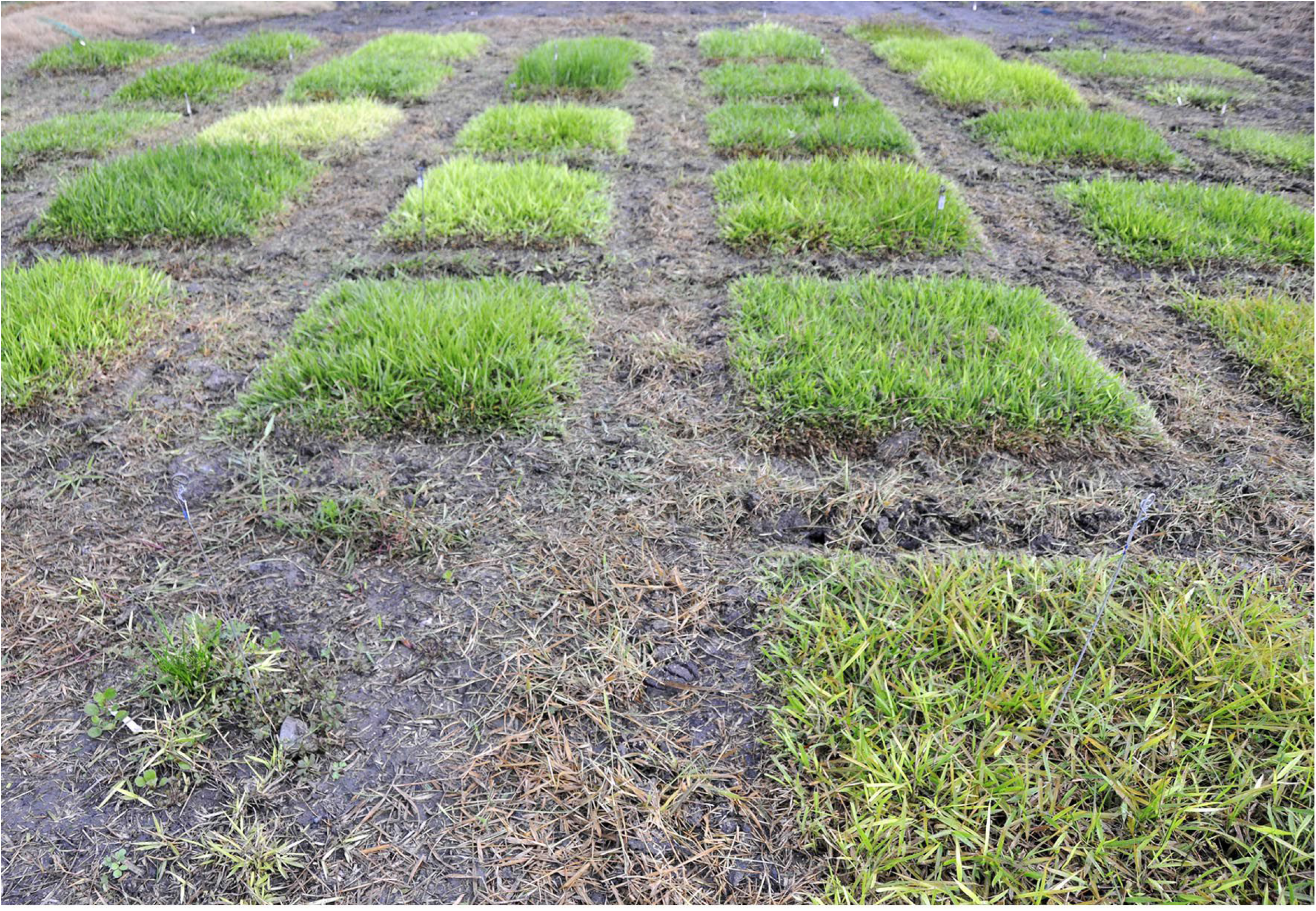
Field plots of *Urochloa* tropical forage grasses in CIAT, Colombia.

### 2.2. Collection and preservation of plant material for flow cytometry

1. Leaf fragments of approximately 1g fresh weight were harvested in the field, folded into permeable manila seed storage envelopes (80gsm) and kept in a sealed plastic bag on wet ice. Young leaves from typical vigorous specimens, representative of the population in each plot, were selected. Insect damaged and discolored plants were avoided.
2. The envelopes were then stored in a sealed desiccators at ambient pressure, or a airtight plastic box (as used for sandwiches or larger sizes), at room temperature with a thick layer of self-indicating silica gel (a granular material with c. 3 to 5mm irregular beads; e.g. Type III Sigma-Aldrich, S7625; or self-indicating mixed with non-indicating silica gel; cheaply available from online marketplaces). The silica gel was changed daily until it did not change colour, which was after approximately 4-5 days. 250g of silica gel was used for 30 leaf samples.
3. Multiple samples in the paper envelopes are then transferred to sealed plastic bags with a small amount of silica gel. If there is any question of insect contamination of leaf collections, the plastic bags can be frozen (−20C, 48hr).
4. The plastic bags with envelopes of dried leaves and silica gel, can be shipped under ambient conditions to the University of Leicester, UK (with appropriate export and import documentation, here under “Section IV: Cut flowers, foliage and vegetables” and “Section III: Seeds for planting” of the UK “Import requirements for plants, plant produce and products”). The sealed bags, after inspection and replacement of silica gel if required, are then stored in 4°C in plastic boxes containing silica gel until flow cytometry analysis.
5. The seeds received from VIR, USDA and CIAT were germinated in a tropical greenhouse (25°C), and leaf samples were collected from plants, and dried and preserve in the same way as those collected in the field in Colombia, and then used as standards for flow cytometry analysis.

### 2.3. Flow cytometry protocol

For ploidy measurement, *Panicum miliaceum* Mil69 (2*n*=4*x*=36; Hunt *et al*., 2014) was used as a first standard to recognize ploidy of some accessions of studied plants. Subsequent internal standards were then included in the analyzes, and their number of chromosomes was confirmed microscopically. Cell nuclei suspension from dehydrated leaf tissues were prepared for flow cytometric analysis according to Doležel *et al*. (2007) with minor modifications.

1. 500 mg of dried leaf of each accession were chopped with a sharp razor blade in a 55×15mm polystyrene Petri dish with ice cold 1 mL nucleus-isolation buffer. Much smaller amounts of leaf material (e.g. 100 mg) did not give suitable nuclear suspensions. We used double edge stainless razor blades (Astra™ Superior Platinum), allocating one razor edge per one studied accession. For safe holding of the razor blade while chopping, a rubber grip was used. Single-edge razor blades are not suitable as they are too thick and not sharp enough. There is variation between different makes of double-edge razor blades: the most widely available Gillette blades can be used but are not as good as some other makes.
2. Three different standard buffers were evaluated, as showed in **Table 1**. Buffers were supplemented with 15mM β-mercaptoethanol and 1% PVP-40 (polyvinylpyrrolidone-40) and the effect of these chemicals on reducing negative effect of cytosolic and phenolic compounds was tested.
3. After finely chopping the material in the buffer, the nuclei suspension was passed through a 50 μm mesh nylon filter (CellTrics, Partec) into the 12×75 mm round-bottom polystyrene flow cytometry tubes (Falcon^®^ with caps preventing cross-contamination, but any other 5mL flow cytometry tubes can be used), and placed on ice.
4. The nuclei suspension was then supplemented with propidium iodide (PI, final concentration 50 μg mL^−1^; solution in deionized water, passed through a 0.22-mm filter), and ribonuclease A (final concentration 50 μg mL^−1^) to prevent staining of double-stranded RNA, and mixed gently using vortex.
5. Samples were incubated at least 10 min (and up to 2 hours) on ice in darkness, and then were analysed in an Accuri C6 Flow Cytometer (Becton Dickinson), equipped with a 20-mW laser illumination operating at 488 nm; however much simpler instruments (e.g. Partec) are sufficient to measure DNA content.
6. The histograms (FSC-A vs SSC-A, FL1-A vs FL2-A, FL3-A vs FL2-A, and an univariate histogram of FL2-A) were acquired using the CFlow^®^ Plus software set up according to Galbraith and Lambert (2012); when use other instruments, follow manufacturer’s instructions for appropriate setting. Here, the following filter configurations were used: FL-1 - a 530/14-nm bandpass filter; FL-2 - a 585/20-nm bandpass filter; and FL-3 - a 670-nm longpass filter. The primary threshold was set to channel 10,000 on FSC-A to gate out debris and noise from nuclei suspension. The secondary threshold was set at 1,000 for FL-2. Polygonal gating tool was used to draw a region on the FSC-A vs SSC-A plot, and a line-shaped cluster of dots showing PI-stained nuclei on the biparametric dot plot of FL2-A vs FL3-A. Based on this gating, G_0_/G_1_ and G_2_ peaks appeared in an univariate histogram of FL2-A.
7. The relative fluorescence intensity of PI-stained nuclei (FL), and the coefficient of variation (CV) of the G_0_/G_1_ peak to estimate nuclei integrity and variation in DNA staining were evaluated in each sample by placing regions of identification across the peak to export values.
8. Ploidy of studied plants were determined by comparing the PI fluorescence intensities of samples to that of standards.

### 2.4. Microscopy and validation of chromosome numbers

For chromosome number calculation of standards we used modified protocol of Schwarzacher and Heslop-Harrison (2000).

1. *Urochloa* seeds, like many other tropical grasses, did not germinate in Petri dishes. The seeds were germinated in a 25°C greenhouse, in a 15×15cm plastic pots containing Levington F2 + S soil.
2. Root tips were collected from plants cultivated in a greenhouse, treated with α-bromonaphthalene at room temperature for 2 h, and 4°C for 4 h, and fixed in absolute ethyl alcohol:acetic acid solution, 3: 1.
3. The root tips were washed in enzyme buffer (10mM citric acid/sodium citrate) for 15 min, and then they underwent enzymatic maceration in 20U/ml 2 cellulase (e.g. Sigma C1184), 10U/ml ‘Onozuka’ RS cellulase and 20U/ml pectinase (e.g. 3 Sigma P4716 from *Aspergillus niger*; solution in 40% glycerol) in 10mM enzyme buffer for 60 min at 37°C.
4. Digested root tips were squashed in 60% acetic acid. Cover slips were removed after freezing with dry ice.
5. Air-dried slides were counterstained with DAPI (4’,6-diamidino-2-phenylindole, 2 μg/mL) in antifade solution (Citifluor, Vectashield, Slowfade or any other commercial antifading reagents for fluorescence microscopy), which prevents the permanent loss of fluorescence due to prolonged exposure to high intensity light sources.
6. Slides were analyzed with an epifluorescence microscope with appropriate UV illumination, filters and camera (Nikon Eclipse 80i; DS-QiMc monochromatic camera, and NIS-Elements v.2.34 software, Nikon, Tokyo, Japan). The number of chromosomes were counted.

## 3. Results

### 3.1. Optimization of flow cytometry assay for dried leaves of Urochloa

#### 3.1.1. Flow cytometry troubleshooting

Nuclei isolated from properly collected, dried and well-preserved leaf samples, as explained in materials and methods, give histograms showing peaks from cells at different stages of the cell cycle: higher G_0_/G_1_ (DNA in nuclei is unreplicated, and may come from differentiated cells) and lower G_2_ (DNA replicated) peaks. Sometimes more peaks are observed, like three gradually declined peaks on **Fig. 2A**, indicating endoreduplication process. Use of older leaf collections may give less marked, flatter, wider or additional peaks on the histogram (**Fig. 2B**). Fresh and dried leaves should give the similar position of peaks, as shown on **Fig. 2C** and 2**D** for comparison, however, the number of isolated nuclei from dried leaves may be smaller due to sample degradation. Ideally, several samples of one accession should be run as the position of the peak on the histogram may vary slightly between plants. It is extremely important to use standard double-edge razor blades as those with single edge are not very sharp, crushing rather than chopping the tissue, resulting in thick and short peaks on histograms (**Fig. 2E**); it is important to chop rather than slice the leaves. In order to get better results and remove debris and noise from histograms, gating is recommended. In the example of **Fig. 2F**, nuclei of interest were being selected (gated) on the FSC-A vs SSC-A and FL2-A vs FL3-A plots (as explained in M & M), resulting in sharper peaks of G_0_/G_1_ and G_2_ and lower background on univariate histogram of FL2-A, in comparison to **Fig. 2G** where gating tools were not applied.

**Figure 2.**
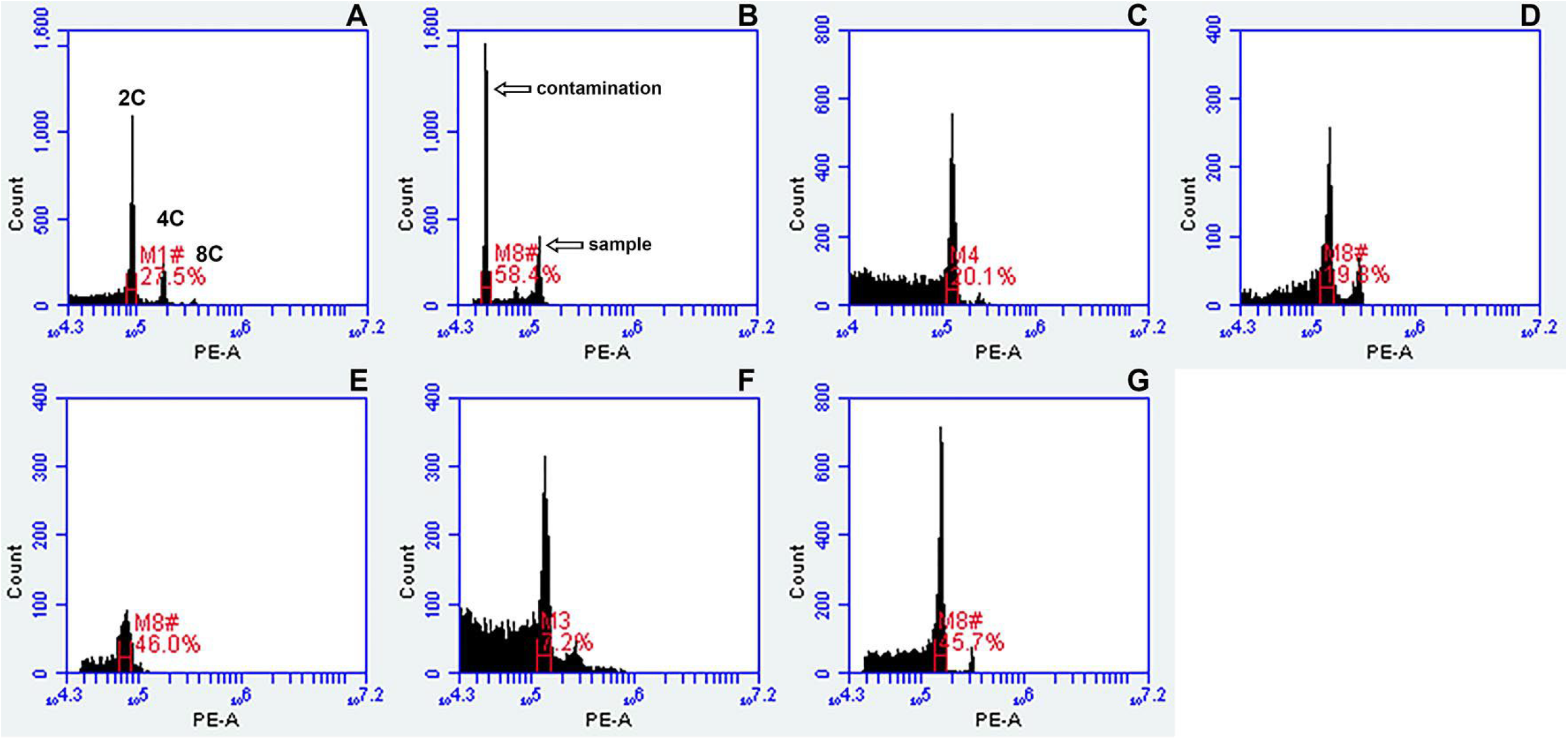
Optimization of flow cytometry assay for dried leaf samples. (A) Three gradually declined peaks of diploid *U. decumbens* CIAT 26185 indicating endoreduplication; (B) Additional high peak on histogram of tetraploid *U. maxima* CIAT 16055, indicating contamination of leaf sample; (C) Fresh and (D) dried leaf samples of *Panicum miliaceum* showing the same position of peaks on histograms, but slight differences in number of nuclei and CV; (E) Histogram of leaf sample chopped with single-edge razor blade; (F) Histogram of dried leaf sample of *Panicum miliaceum* with no gating tools applied; (G) Histogram of dried leaf sample of *Panicum miliaceum* where gating tools were applied, giving sharp peaks and low background.

#### 3.1.2. Buffers

Three different standard isolation buffers (**Table 1**) were tested to isolate nuclei from dehydrated *Urochloa* leaves. No peaks (**Fig. 3A**) or very low peaks were obtained analyzing samples of nuclei isolated using Galbraith’s buffer (Galbraith *et al*., 1983) which is optimized for fresh material. Small numbers of nuclei were isolated using Otto’s buffer (Otto, 1992) giving histograms with increased level of background and high CVs (**Fig. 3B**). Supplementation of Otto’s buffer with β-mercaptoethanol only slightly increased the peak resolution (**Fig. 3C**). Well-defined histograms with acceptable CV values and reasonable number of nuclei (**Fig. 3D,E,F**) were obtained using Partec buffer (de Laat *et al*., 1987). The sharpest peaks were yielded after supplementation of this buffer with 15mM β-mercaptoethanol and 1% PVP-40 (**Fig. 3F**).

**Figure 3.**
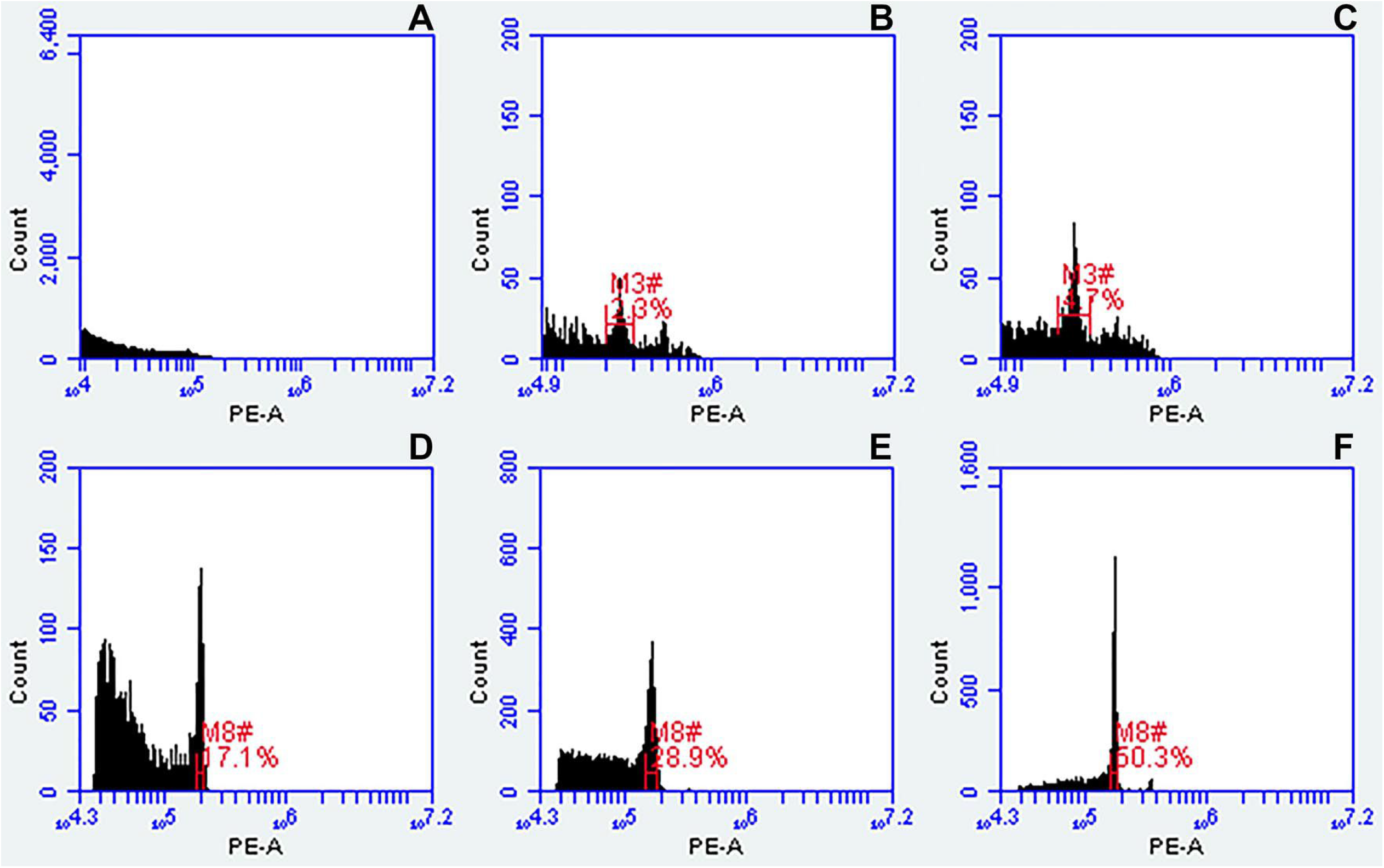
Comparison of three different standard buffers for nuclei isolation from dried leaves of tetraploid *Urochloa* accessions, and their effect on DNA cell cycle histogram quality. (A) Galbraith’s buffer; (B) Otto’s buffer; (C) Otto’s buffer supplemented with β-mercaptoethanol; (D) Partec buffer; (E) Partec buffer supplemented with β-mercaptoethaπol; (F) Partec buffer supplemented with 15mM β-mercaptoethanol and 1% PVP-40. Regions of identification (red) were placed across the peaks to export values representing peak positions and CVs.

#### 3.1.3. Standards used for flow cytometry analysis

The procedure of counting ploidy from *Urochloa* dried leaf specimens by flow cytometry was optimized by choosing appropriate buffer composition (**Table 1, Fig. 3**), drying and preservation of plant samples (protocol in M&M, **Fig. 2**), chopping technique (**Fig. 2**) and eleven different standards (**Table 2, Fig. 4**). *Panicum miliaceum* (2*n*=2*x*=36) was used as a first standard to recognize accessions for which the level of ploidy was certain. The seeds of these accessions were obtained from CIAT and USDA, germinated in a greenhouse, and the ploidy of plants was validated by preparing mitotic slides and counting chromosomes microscopically (**Fig. 4**). These samples were then used as internal standards, and their mean peak indices were given in **Table 2**. The position of peaks of *Urochloa humidicola* CIAT 16867 on the histogram (**Fig. 4F**) suggested that this accession was most likely to be heptaploid, but chromosome counting revealed it to be aneuploid with 2*n*=8*x*+2 or 9*x*-4=50.

**Figure 4.**
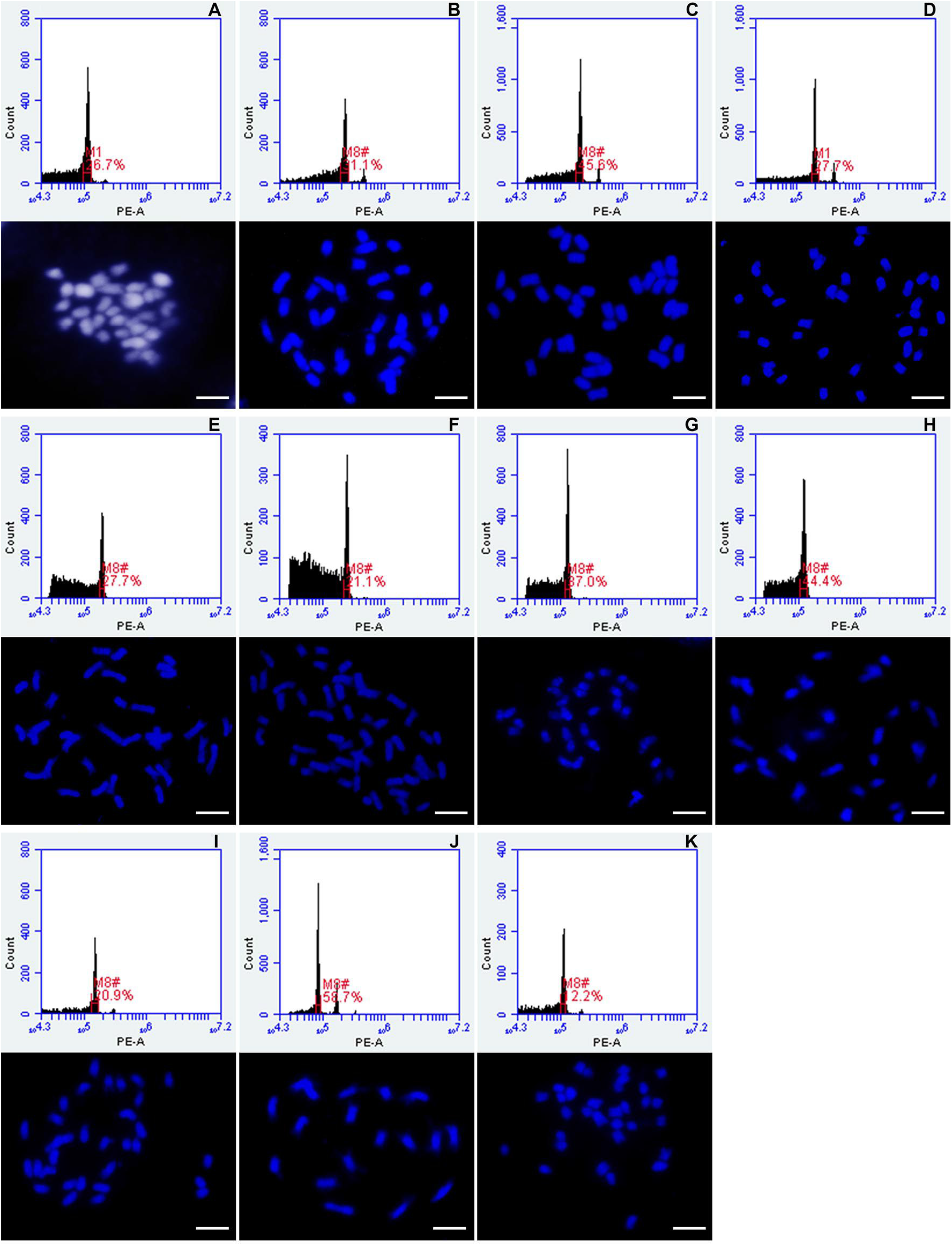
Histograms of relative fluorescence intensities showing ploidy levels and the corresponding chromosome numbers of different genotypes used as standards for flow cytometry analysis of *Urochloa* germplasm collection. Standard peak means in Table 2. (A) *Panicum miliaceum* Mil69 (2*n*=2*x*=36); (B) *Urochloa brizantha* PI292187 (2*n*=4*x*=36); (C) *Urochloa decumbens* CIAT 664 (2*n*=4*x*=36); (D) *Urochloa decumbens* CIAT 6370 (2*n*=4*x*=36); (E) *Urochloa humidicola* CIAT 26151 (2*n*=6*x*=36); (F) *Urochloa humidicola* CIAT 16867 (2*n*=8*x*+2 or 9*x*-4=50); (G) *Urochloa maxima* CIAT 6171 (2*n*=4*x*=32); (H) *Urochloa maxima* CIAT 16004 (2*n*=4*x*=32); (I) *Urochloa maxima* PI 284156 (2*n*=4*x*=32); (J) *Urochloa ruziziensis* CIAT 6419 (2*n*=2*x*=18); (K) *Urochloa* sp. PI 657653 (2*n*=4*x*=32). Regions of identification seen on plots (red) were placed across the peaks to export values representing peak positions and CVs. Scale bars = 5μm.

**Table 2.** Standards used for flow cytometry analysis of *Urochloa* germplasm collection. Chromosome numbers were counted microscopically.

| Standard | Accession number | Number of chromosomes | Standard mean peak |
| --- | --- | --- | --- |
| *Panicum miliaceum* | Mil69 | 2n=4x=36 | 111,925.27 |
| *Urochloa brizantha* | PI 292187 | 2n=4x=36 | 225,075.73 |
| *Urochloa decumbens* | CIAT 664 | 2n=4x=36 | 205,253.15 |
| *Urochloa decumbens* | CIAT 6370 | 2n=4x=36 | 193,675.46 |
| *Urochloa humidicola* | CIAT 26151 | 2n=6x=36 | 197,353.12 |
| *Urochloa humidicola* | CIAT 16867 | 2n=8x+2 or 9x-4=50 | 252,917.76 |
| *Urochloa maxima* | CIAT 6171 | 2n=4x=32 | 130,912.51 |
| *Urochloa maxima* | CIAT 16004 | 2n=4x=32 | 119,920.97 |
| *Urochloa maxima* | PI 284156 | 2n=4x=32 | 148,800.27 |
| *Urochloa ruziziensis* | CIAT 6419 | 2n=2x=18 | 82,708.77 |
| *Urochloa* sp. | PI 657653 | 2n=4x=32 | 110,639.72 |

### 3.2. Ploidy measurement of *Urochloa* and related species

DNA content of 353 accessions of *Urochloa* and related species from CIAT and USDA germplasm collection were measured using flow cytometry of imported dried leaf materials using the optimized technique giving very sharp peaks. Values representing peak positions and CVs were exported and are given in **Supplementary Data Table S1**, and summarized in **Table 3**. CV values were slightly increased comparing to the fresh leaf specimens of *Panicum miliaceum* (approximately 2,5%). A coefficient of variation of less than 5% is desirable, but analysis of older leaves often gives broader peaks with a still usable CV between 5% and 10%. Where possible, several leaf samples for one accession were measured enabling comparison of mean peak positions between different plants of the same accession. In general, these values did not differ significantly from plant to plant, proving the established method. The position of mean peak samples were compared to that of the eleven standards (**Table 2**). For each level of ploidy of the individual species, a range of mean peak indices have been established (**Table 3**), and these ranges for the most numerous species were shown on **Fig. 5**.

**Figure 5.**
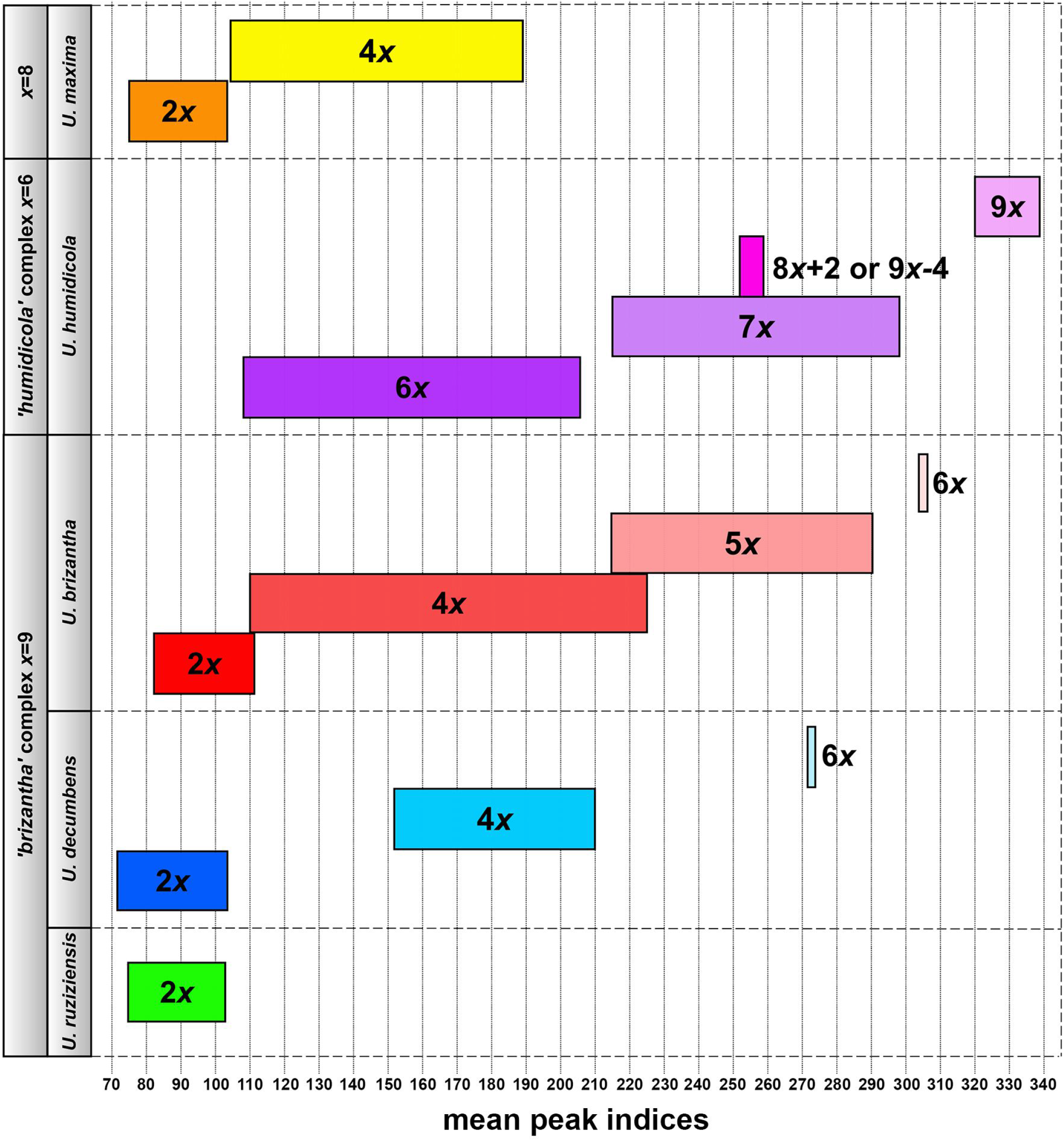
Ranges of mean peak indices for different ploidy levels of the most numerous species (‘*brizantha*’ agamic complex: *U. ruziziensis, U. decumbens, U. brizantha; ‘humidicola*’ agamic complex: *U. humidicola; U. maxima*) in CIAT germplasm collection.

**Table 3.**
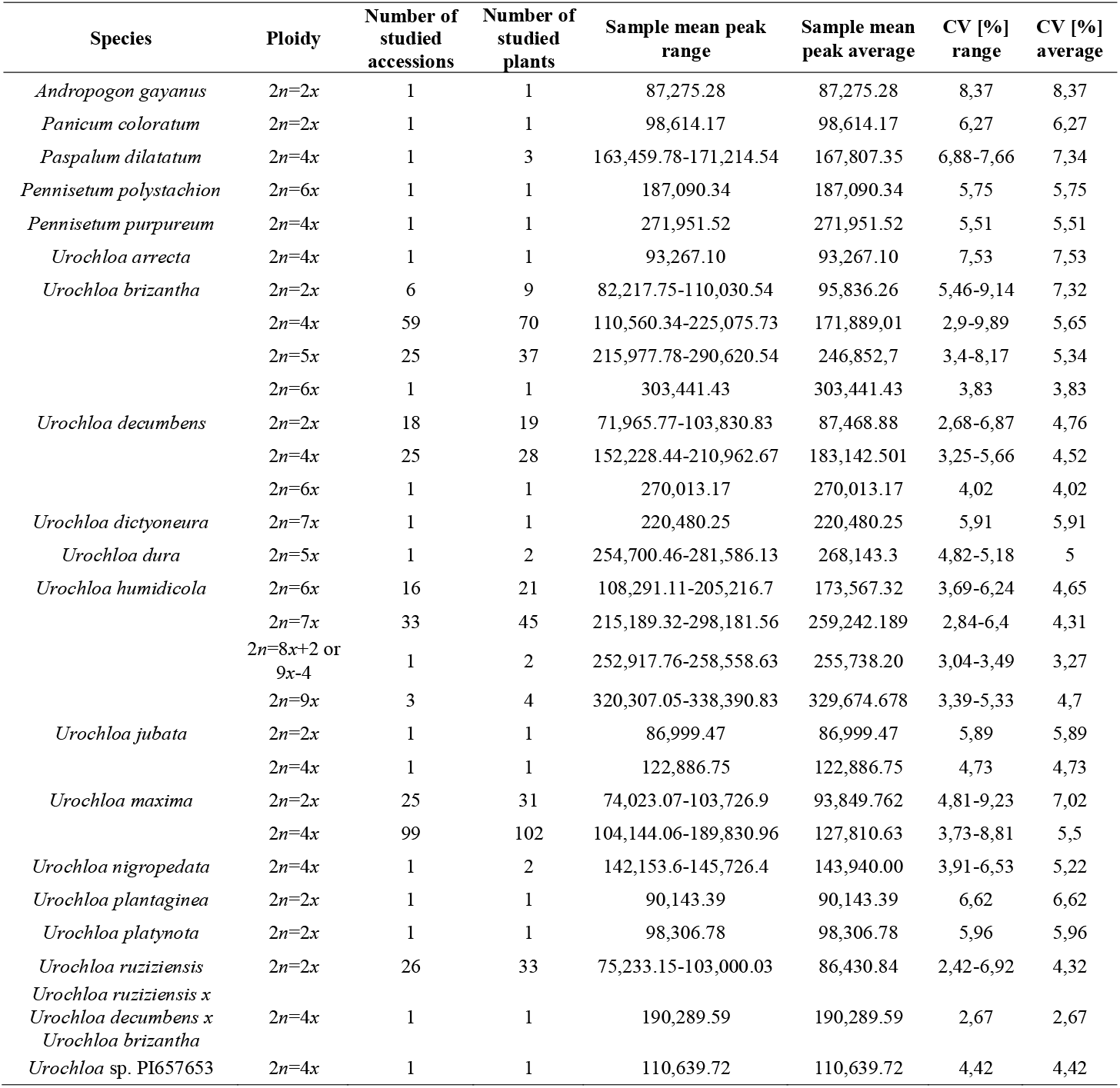
Variation in sample mean peak indices between species, for the different ploidy levels determined.

#### 3.2.1. ‘*brizantha*’ agamic complex

Three species belonging to the ‘*brizantha*’ agamic complex: *Urochloa ruziziensis, U. decumbens* and *U. brizantha* have a basic chromosome number *x*=9. All accessions of *U. ruziziensis* studied here were diploid (**Supplementary Data Table S1**), showing similar range and average of mean peak indices to that of diploid *U. decumbens* (see **Table 3** and **Fig. 5**). Within both species there are single samples showing higher mean peak indices than the others. *U. decumbens* accessions differ in their ploidy levels, showing diploids, tetraploids, and hexaploid (**Supplementary Data Table S1**), that can be clearly distinguishable using flow cytometry, because the ranges of mean peak indices for each ploidy level did not overlap. This results contrasts with *U. brizantha*, where the sample mean peak ranges of diploids, tetraploids, pentaploids and hexaploids overlapped, meaning that ploidy levels of this species are not so obvious (see **Fig. 5)**. This is particularly evident when looking at the differences in index values between samples of the same accession (see **Supplementary Data Table S1**).

#### 3.2.2. ‘*humidicola*’ agamic complex

Two polyploid species with basic chromosome number *x*=6 were assigned to the ‘*humidicola*’ agamic complex: *U. humidicola* and *U. dictyoneura*. Three different ploidy levels were recognized in the *U. humidicola*: hexaploid, heptaploid, and nonaploid (**Supplementary Data Table S1**). *U. dictyoneura* accession used in our studies seemed to be heptaploid. In general, each ploidy level of *U. humidicola* has its own range of mean peak values (**Fig. 5**), however due to confirmed aneuploidy within species (*U. humidicola* CIAT 16867 with 2*n*=8*x*+2 or 9*x*-4=50), additional validation, e.g. counting chromosome numbers, seems to be needed.

#### 3.2.3. Urochloa maxima

Two ploidy levels were recognized in *U. maxima*. Some diploid and tetraploid accessions showed similar values of mean peaks (**Fig. 5**). Those samples that have extreme results and peaks well beyond those of the reference internal standards, should have their chromosomes counted.

#### 3.2.4. Related species

Several tropical grass species with potential for improvement and wider use as forages have been studied here, including other cultivated and wild *Urochloa* species, as well as *Paspalum, Panicum, Pennisetum*, and *Andropogon*, showing different basic chromosome numbers. In most cases, our internal standards were useful to establish ploidy levels of studied species. However, for *Pennisetum polystachion* and *P. purpureum* with basic chromosome numbers *x*=9 and *x*=7, respectively, we had to use the literature data on possible ploidy levels observed for these species due to the higher genome size comparing to the internal standards belonging to ‘*brizantha*’ and ‘*humidicola*’ complexes (Martel *et al*., 1997; Campos *et al*., 2009; dos Reis *et al*., 2014).

## 4. Discussion

### 4.1 Flow cytometry as a standard technology

Flow cytometry has become the standard technology for measuring the ploidy and genome sizes of plants (Doležel and Bartoš, 2005), allowing the measurement of hundreds of samples, even genomically diverse species, in a relatively short time. In most cases, freshly collected, field- or garden-grown leaf material is used, with a small number of accessions. We have optimized the methods for sampling, drying, storage, transport and preservation of tropical forage grasses to use some time later with a robust flow cytometry protocol for measurement of ploidy (**Figs 2, 3**; **Table 1**). We show the utility in a relatively large and diverse germplasm collection (353 accessions) of the tropical forage grass genus *Urochloa (Brachiaria*) (**Tables 2, 3; Supplementary Data Table 1**). The method allows wider field and geographical sampling of plants when fresh leaf tissues cannot be examined shortly after harvesting (Wang and Yang, 2016). Integration of ploidy levels and agronomic traits, especially those related to resistance and tolerance to pest and diseases, is important to define a breeding strategy to exploit germplasm with diverse ploidy levels (Alves *et al*., 2013; Barrios *et al*., 2013; Matias *et al*., 2016). Where collections have various ploidies, flow cytometry can help the verification of samples from field collections, where mislabelling, or spread of incorrect seed or plants in vegetative plots may lead to replacement of one accession with another over decades. Comparison of similar accessions numbers from Brazil and Colombia detects some such differences. Polyploidy promotes genome diversification and gives plasticity to species (Soltis and Soltis, 1993), thus it is pertinent to examine ploidy of as many accessions as possible in order to choose those suitable for crossbreeding. For research purposes, sampling and screening the large germplasm collections provides additional characters and help to better estimate genome relationships between species within large plant complexes, such as *Urochloa* (González and Morton, 2005) and hence help reconstruct phylogenies, particularly those where reticulate evolution of polyploid taxa is found. Flow cytometry and the measurement of nuclear DNA contents has other applications not considered here, in particular for determination of cell cycle times (Francis *et al*., 2008), and examining differentiation of cells through endopolyploidy (Bhosale *et al*., 2018). While flow sorting of chromosomes would require living materials, it is likely that dried leaf material may be used to study genome size differentiation patterns in the context of cell cycle times and endopolyploidy.

### 4.2. Choice of approaches of flow cytometry

For flow cytometry-based estimation of DNA content, almost every fluorescence-based flow cytometer can be used, but the filter set should be compatible with the spectral properties of fluorescent dyes (Doležel *et al*., 2007). Propidium iodide (PI) is one of the most widely used fluorescence reagent in flow cytometry binding to DNA by intercalating between DNA bases (rather than then major or minor groove), and showing no AT or GC preference (**Fig. 6**). Its fluorescence with green-light excitation is enhanced some 20-fold when bound to DNA compared to in solution. The emission maximum depends on the solvent, and in the aqueous solution used for nuclear isolation, the maximum is 636 nm (red) (Samanta *et al*., 2012). PI shows intermolecular proton transfer reaction in solvent; it interacts with SDS (sodium dodecyl sulphate), so this widely used detergent cannot be a component of a nuclei isolation buffer for flow cytometry. Propidium iodide also binds to RNA, showing enhanced fluorescence (with a slightly different fluorescent colour), so for nuclear staining, RNase needs to be added to the buffer. In practice, the concentration of PI and RNase in the buffer is important, and peaks broaden (higher CV) when they are too high or too low. Other components include Tris as buffer, NaCl (85mM) to maintain nuclear integrity, β-mercaptoethanol as antioxidant, and PVP-40 (polyvinylpyrrolidone-40) to bind polyphenols and anthocyanins, scavenge other polar molecules and deactivate proteins from the plant cells; and Triton X-100 as a detergent to aid buffer penetration.

**Figure 6.**
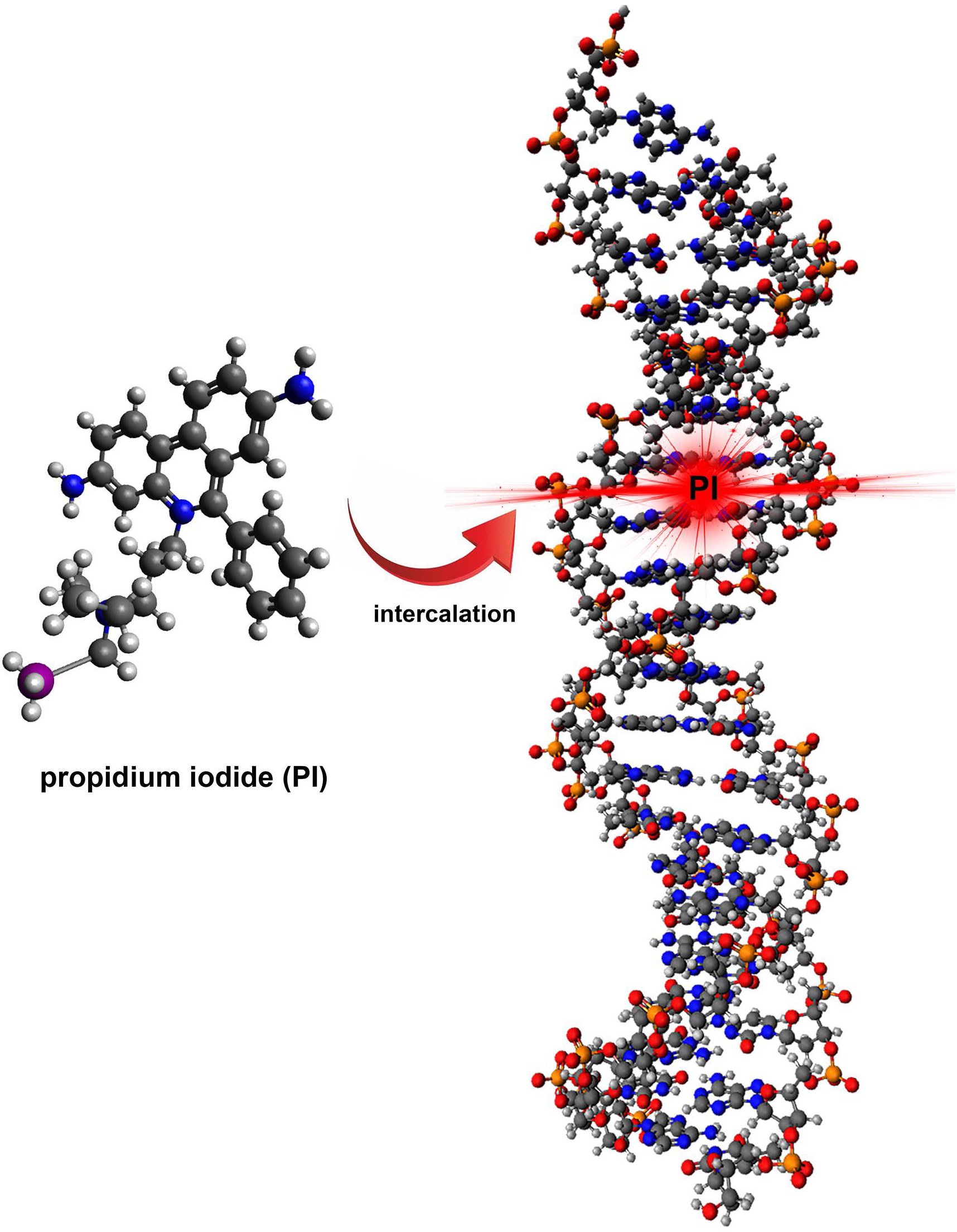
Diagram showing intercalation of propidium iodide molecule between DNA bases. Avogadro program (Hanwell *et al*., 2012) was used to create a molecule of propidium iodide, and a Watson-Crick duplex using the sequence AATAACTCCCACATGTCCAT.

The chopping with a very sharp razor blade is a critical part of the technique. If the blade is wrongly used with slicing motion, or has a dull edge, the nuclei are sheared and the peaks become very broad.

### 4.3. *Urochloa* germplasm findings

Here, we verified ploidy of 353 accessions of *Urochloa* and related species, which represent a significant proportion of CIAT germplasm resources. However, determining the ploidy levels of grass group showing both apomictic and sexual mode of reproduction, like *Urochloa*, can become a challenge and requires the use of appropriate standards of known ploidy and number of chromosomes (Krahulcová and Rotreklová, 2010). For *Urochloa* grass complex, different internal standards were needed due to the different genome sizes within and between agamic complexes and species, and different basic chromosome numbers (**Supplementary Data Table 1**). The average DNA content and genome sizes given as Cx values have been published already for *Urochloa* species (Ishigaki *et al*., 2010; Timbó *et al*., 2014). Most diploid accessions of *U. brizantha* studied here are apomict (Tomaszewska *et al*., 2021), showing larger mean peak indices than sexual diploid accessions of *U. decumbens* and *U. ruziziensis*, proving that the genome size depends on the mode of reproduction (Ishigaki *et al*., 2010), which is an additional challenge for screening diverse germplasm collections. While in diploid and polyploid accessions of *U. decumbens* a small shift in peak position on histogram usually does not compromise reliability of ploidy estimates, attention should be paid to the analysis of *U. brizantha* showing odd ploidy levels, because relative differences in nuclear DNA content between neighboring cytotypes (2*x*, 4*x*, 5*x*, 6*x*) is decreased (see **Fig. 5**); and such a phenomenon is also observed in species with ploidy levels greater than 6*x* (Doležel *et al*., 2007). A more precise examination of *U. humidicola* is also required due to confirmed aneuploidy (see **Figs 4F** and **5**; Moraes *et al*., 2019; da Rocha *et al*., 2019), odd ploidy levels (Boldrini *et al*., 2009b), and unrecognized diploid ancestors (Boldrini *et al*., 2009a).

*Urochloa* tropical forage grasses and related genera studied here, including *Andropogon, Pennisetum, Paspalum*, and *Panicum* have a great potential for sustainable agriculture and intensive grazing management of cover crops. Some of them are included in the current breeding programs at CIAT and EMBRAPA, now mainly focused on crossing tetraploids within ‘*brizantha*’ and ‘*humidicola*’ agamic complexes and *Urochloa maxima* (Triviño *et al*., 2017). These tropical forage grass group is genomically complex (Tomaszewska *et al*., 2021), having species recognized as being very variable in number of chromosomes, and ploidy levels which is the result of apomictic reproduction, and reflecting the genetic diversity present in a given population (Jank *et al*., 2011). The ploidy levels of some *Urochloa* accessions have been previously measured (Penteado *et al*., 2000; Jungmann, 2009; Jungmann *et al*., 2010; Nitthaisong *et al*., 2016; Triviño *et al*., 2017), but some data vary between papers and reports (Tomaszewska *et al*., 2021): thus values may requires checking for a particular accession name.

## Supplementary Materials

**Supplementary Data Table S1** List of accessions used in the study, their mean peak indices and coefficient of variations (CVs) of the G_0_/G_1_ peaks.

## Acknowledgments

This work was supported under the RCUK-CIAT Newton-Caldas Initiative “Exploiting biodiversity in *Brachiaria* and *Panicum* tropical forage grasses using genetics to improve livelihoods and sustainability”, with funding from UK’s Official Development Assistance Newton Fund awarded by UK Biotechnology and Biological Sciences Research Council (BB/R022828/1). PT has received further support from the European Union’s Horizon 2020 research and innovation programme under the Marie Sklodowska-Curie grant agreements No 844564 and No 101006417.

We would like to thank Dr Jennifer Hincks from The Centre for Core Biotechnology Services, Flow Cytometry Facility at University of Leicester. We are grateful to USDA-Germplasm Resources Information Network (GRIN), and Vavilov Research Institute (VIR, St Petersburg, Russia) for their generous provision of seeds. Germplasm held in the CIAT collections is available on request (http://genebank.ciat.cgiar.org).

## Author Contributions

Writing—original draft: PT; Conceptualization: PT, TP, VC and PH-H; Investigation - flow cytometry analysis, chromosome preparation and counting: PT; Sampling: TP, LM; Methodology: PT, TP; Funding acquisition: RM, JV and PH-H; Supervision: PH-H;

Writing—review & editing: PT, TP, LM, RM, VC, JV, TS and PH-H. All authors have read and agreed to the published version of the manuscript.

